# Can AI reproduce observed chemical diversity?

**DOI:** 10.1101/292177

**Authors:** Mostapha Benhenda

**Author notes:** Correspondence Startcrowd, online AI lab, Vyborska Street, 80/17 apt. 63, Kiev, Ukraine Full list of author information is available at the end of the article.

## Abstract

Generating diverse molecules with desired chemical properties is important for drug discovery. The use of generative neural networks is promising for this task. To facilitate evaluation of generative models, this paper introduces a metric of internal chemical diversity, and raises the following challenge: can a nontrivial AI model reproduce observed internal diversity for desired molecules? To illustrate this metric, a mini-benchmark is performed with two generative models: a Reinforcement Learning model and the recently introduced ORGAN. The aim of this paper is to encourage research about internal diversity metrics.

## 1 Introduction

Drug discovery is like finding a needle in a haysack. The chemical space of potential drugs contains more than 10^60^ molecules. Moreover, testing a drug in a medical setting is time-consuming and expensive. Getting a drug to market can take up to 10 years and cost $2.6 billion [1]. In this context, computer-based methods are increasingly employed to accelerate drug discovery and reduce development costs.

In particular, there is a growing interest in AI-based generative models. Their goal is to generate new lead compounds *in silico*, such that their medical and chemical properties are predicted in advance. Examples of this approach include Variational Auto-Encoders [2], Adversarial Auto-Encoders [3, 4], Recurrent Neural Networks and Reinforcement Learning [5, 6, 7], eventually in combination with Sequential Generative Adversarial Networks [8, 9].

However, research in this field often remains at the exploratory stage: generated samples are sometimes evaluated only visually, or with respect to metrics that are not the most relevant for the actual drug discovery process.

Rigorous evaluation would be particularly welcome regarding the **internal chemical diversity** of the generated samples. Generating a chemically diverse stream of molecules is important, because drug candidates can fail in many unexpected ways, later in the drug discovery pipeline.

Based on visual inspection, [5, p. 8] reports that their Reinforcement Learning (RL) generative model tends to produce simplistic molecules. On the other hand, [8, p.6, p.8] argues that their Objective-Reinforced Generative Adversarial Network (ORGAN) generates less repetitive and less simplistic samples than RL. However, their argument is also based on visual inspection and therefore, it remains subjective: our own visual inspection of the ORGAN-generated samples (available on the ORGAN Github:

https://github.com/gablg1/ORGAN/tree/master/results/mol_results) rather suggests that ORGAN produces molecules as repetitive and as simplistic as RL.

In this paper, we introduce a metric that quantifies the *internal chemical diversity* of the model output. We also submit a challenge:

### Challenge

Is it possible to build a non-trivial generative model, with (part of) its output satisfying a non-trivial chemical property, such that the internal chemical diversity of this output is at least equal to the observed diversity naturally found for the same kind of molecules?

To illustrate this challenge, we compare RL and ORGAN generative models, with respect to the following chemical properties:

1. Being **active against the dopamine receptor D2**. The dopamine D2 receptor is the main receptor for all antipsychotic drugs (schizophrenia, bipolar disorder...).
2. **Drug-likeness** as defined in [8]. We are interested in this property because we can use experimental results in [8] to facilitate discussion. However, the notion of druglikeness in [8] is different from the notion of Quantitative Estimation of Druglikeness (QED) [10], which is an index measuring different physico-chemical properties facilitating oral drug action.

Here, druglikeness is the arithmetic mean of the solubility (normalized logP), novelty (which equals 1 if the output is outside of the training set, 0.3 if the output is a valid SMILES in the training set, and 0 if the output is not a valid SMILES), synthesizability (normalized synthetic accessibility score [11]) and conciseness (a measure of the difference of the length between the generated SMILES and its canonical representation).

We mention that recently, [9] considers an ORGAN with the QED definition of druglikeness. However, we also performed our own experiments with the QED property, and they did not affect our conclusions.

## 2 The metric of internal chemical diversity

Let *a* and *b* be two molecules, and *m_a_* and *m_b_* be their Morgan fingerprints [12]. Their number of common fingerprints is *|m_a_ ∩ m_b_|* and their total number of fingerprints is *|m_a_ ∪ m_b_|*.

The Tanimoto-similarity *T_s_* between *a* and *b* is defined by:

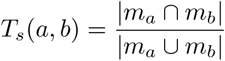

Their Tanimoto-distance is:

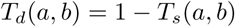

We use rdkit implementation [13] of this distance, with fingerprint size 4.

### 2.1 Internal diversity

We define the *internal diversity I* of a set of molecules *A* of size *|A|* to be the average of the Tanimoto-distance *T_d_* of molecules of *A* with respect to each other. Formally, we have:

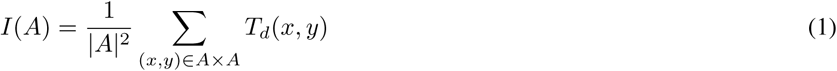

Note that this sum includes self-distances, although their contributions are equal to zero.

For a sufficiently large set *A*, any sufficiently large subset *A^’^ ⊂ A*, sampled with uniform probability, has the same internal diversity as *A*. This property follows from the law of large numbers. We can thus define the internal diversity of a generative model, by computing the internal diversity of a sufficiently large generated sample. This allows to formalize our challenge:

#### Challenge (restatement)

Let *N* be the molecules observed in nature. Is there a nontrivial generative model *G* and a non-trivial chemical property *P* such that:

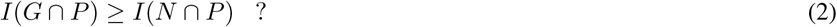

Internal chemical diversity is always smaller than 1 (because the Tanimoto-distance is smaller than 1), and it is usually much smaller. That’s why we prefer this definition to the Tanimoto-variance of a set of molecules *A*, which is:

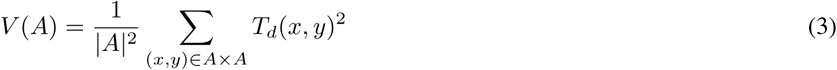

### 2.2 Relation of the internal diversity metric with the previous literature

Internal diversity quantitatively captures the visually observed fact that generated molecules can be repetitive and simplistic [8, 5]. Previous metrics did not allow to do that.

#### 2.2.1 Internal vs. external diversity metric [8, p.5]

Let *A*_1_ be the training set, and *A*_2_ be the generated set. The *external diversity* (called ‘diversity’ in [8, p.5]) is defined by:

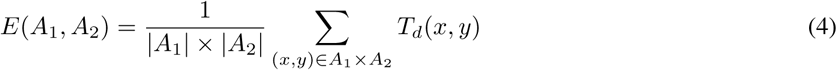

External diversity and internal diversity metrics are different: in our definition we only have one set *A*, the generated set (see equation (1)).

External diversity fails to capture the visually observed fact that generated molecules can be repetitive and simplistic (as observed in [8, 5]): according to table 1 page 5 in [8], (external) diversity is significantly higher for RL than for ORGAN. That’s the opposite of their debatable visual observation.

**Table 1.** Experimental results for probability of D2 activity *P_a_*

| | Prop. Valid SMILES | Avg. $P_a$ | Avg. int. div. | Prop. $P_a > 0.8$ | Avg. ext. div. for $P_a > 0.8$ | Avg. int. div. for $P_a > 0.8$ |
| --- | --- | --- | --- | --- | --- | --- |
| MLE | 0.290406 | 0.009643 | 0.393832 | 0 | $n/a$ | $n/a$ |
| DRD2 | 0.996636 | 0.911519 | 0.089478 | 0.876367 | 0.353919 | 0.081972 |
| RL 30 | 0.379844 | 0.160777 | 0.112864 | 0.018906 | 0.638772 | $8.65864e - 05$ |
| RL 60 | 0.536 | 0.389979 | 0.014994 | 0.078438 | 0.850716 | 0.000775 |
| ORGAN-0.04 30 | 0.425375 | 0.097810 | 0.242544 | 0.013531 | 0.540684 | 0.005826 |
| ORGAN-0.04 60 | 0.604406 | 0.342687 | 0.028563 | 0.100969 | 0.575253 | 0.000170 |
| ORGAN-0.5 60 | 0.264687 | 0.006502 | 0.324884 | 0.000187 | 0.527566 | 0 |

They dismiss RL generator in favor of their ORGAN generator only by their visual observation, but nowhere, their quantitative results support their observation.

On the other hand, our metric will give better results, because it better matches human visual observation of samples: our diversity is slightly lower for RL than for ORGAN.

Why internal diversity metric works better than external diversity metric:

Suppose the chemical space is ℝ^2^ with the Euclidean distance (in place of the Tanimoto-distance). Suppose the training data is located on a circle of radius one centered in the origin. Suppose also a trivial generative model, with all generated samples located on this origin. See figure 1.

**Figure 1:**
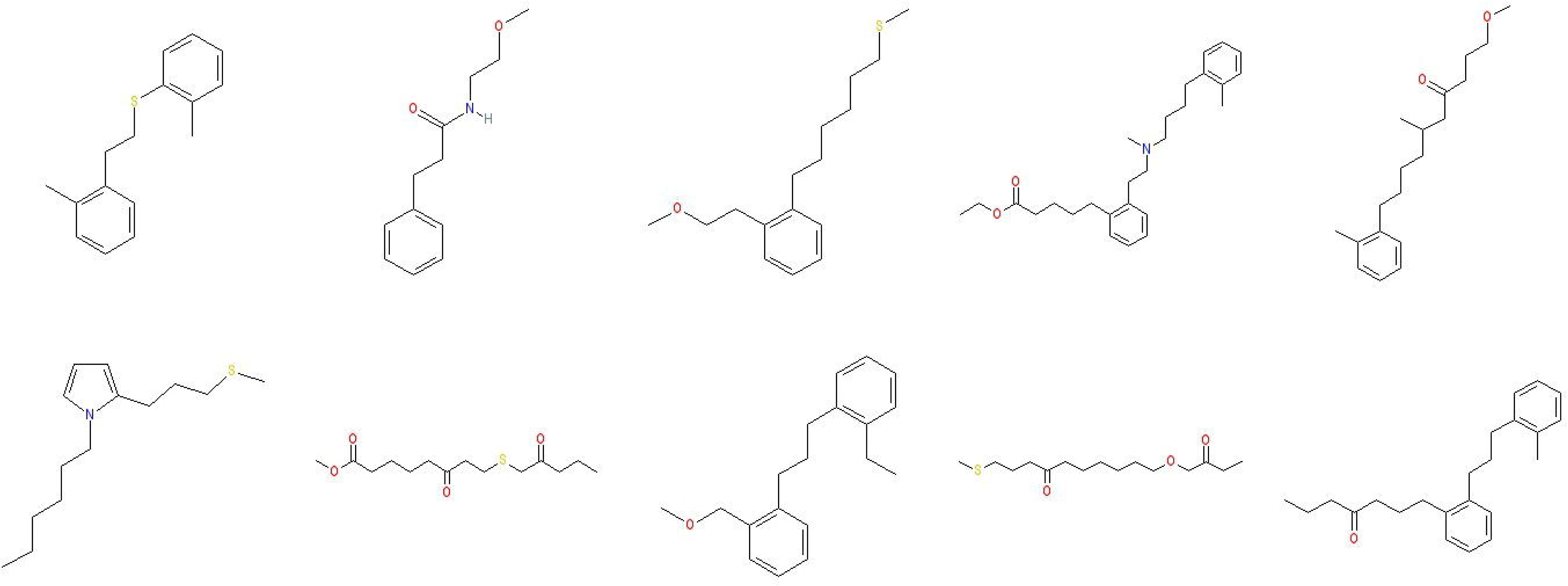
Trivial generative model. Generated samples all coincide on the blue dot, and training data are the red crosses. External diversity= 1, and internal diversity =0

In this setting, the external diversity of the model is equal to 1, because the distance between a generated point and a training point is always equal to 1.

On the other hand, the internal diversity for this generative model is equal to zero, because the distance between two generated samples is always zero.

Contrary to the external diversity metric in [8], our metric can distinguish between this trivial case and a less trivial generative model, where generated points are spread around (figure 2).

**Figure 2:**
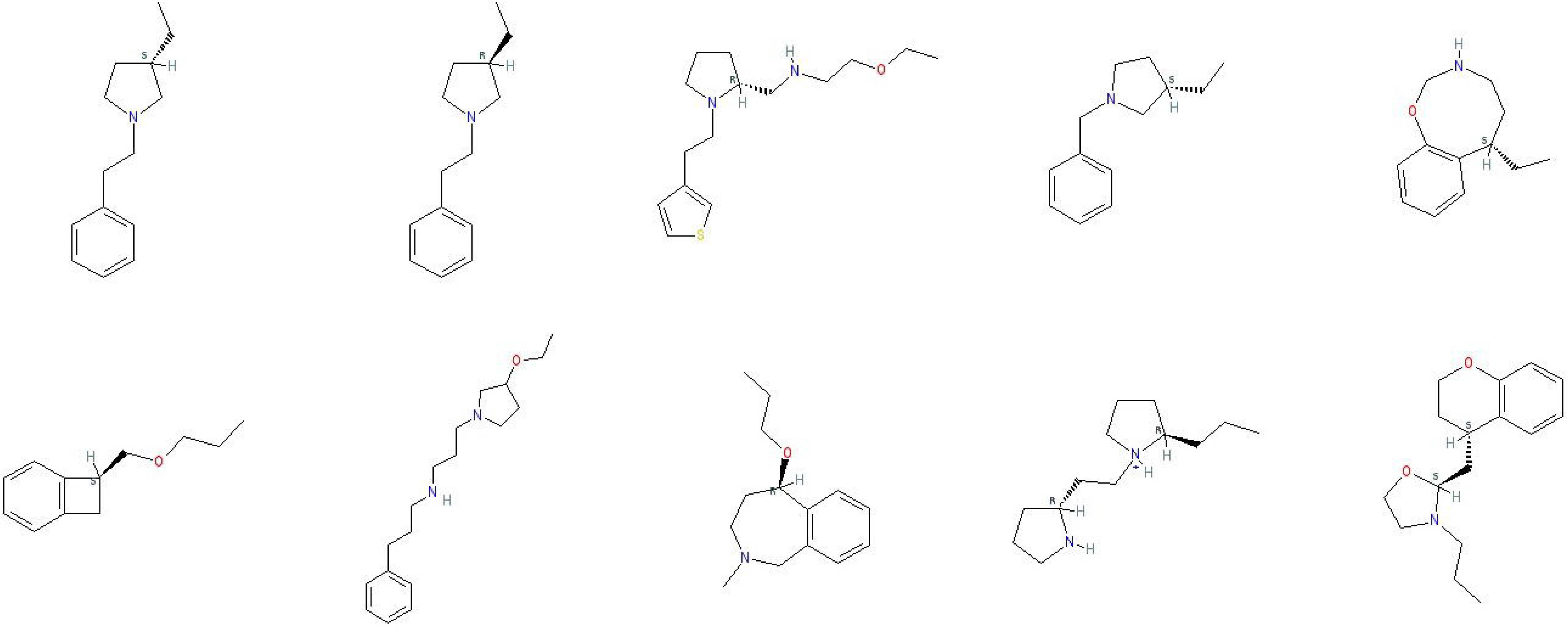
Non-trivial generative model. Generated samples (blue dots) are spread around. Training data are the red crosses.

However, some sort of external diversity metrics is still important, in order to eliminate another kind of trivial generative model, which simply reproduces the training set. There is complementarity between suitable external and internal diversity metrics.

#### 2.2.2 Internal diversity metric vs. Guimaraes et al. [8] novelty metric

In [8, p.4], Guimaraes et al. introduce a *novelty* metric, equal to 1 if the generated molecule is not in the training set, equal to 0.3 if it is inside the training set, and equal to 0 if it is not a valid SMILES.

Our trivial example above would have a high Guimaraes-novelty (all generated molecules are outside of the training set), so Guimaraes-novelty is less useful than our metric for the task of identifying some trivial generators.

#### 2.2.3 Internal diversity metric vs. NN-Tanimoto similarity [6, p. 7 and figures 7 and 12]

In [6, p. 7 and figures 7 and 12], Segler et al. introduce a metric of Nearest Neighbor Tanimoto similarity. Let *m* be a molecule and *A*_1_ be the training set. The Nearest Neighbor Tanimoto-similarity between *m* and *A*_1_ is given by:

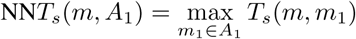

Segler et al. consider the distribution of the Tanimoto-similarities between generated molecules and their nearest neighbor in the training set. They qualitatively discuss the shape of this distribution for their own models, but they do not use this distribution to define a quantitative metric that allows ranking different models.

To extend the work done in [6], various metrics can be derived from this distribution (e.g. variance, Wasserstein distance to the uniform distribution...). This will be interesting for future work, but in any case, these metrics will be more complicated than our metric.

#### 2.2.4 Internal diversity metric vs. NN-Levenshtein distance [6, p. 10 and figure 11]

The same remarks as with the NN-Tanimoto-distance apply. For future work, it will be interesting to replace the Tanimoto distance in our work with the Levenshtein distance.

#### 2.2.5 Internal diversity metric vs. visualizations [6, figures 5 and 8], [14, figures 5, 6 and 9] and [15, figure 2]

Recently, [15, figure 2] used a visualization to claim that the molecules generated by their RL model (similar to [6]) “populate” the chemical space. It is analogous to visualizations in [14, figures 5, 6 and 9], and [6, figures 5 and 8] (except that in this latter case, it is a t-SNE visualization, which is useful, but can also be misleading, see [16]).

The internal diversity metric introduced here is a contribution to give a precise meaning to this expression “populate”. This metric allows to compare different models, from the viewpoint of which one “populates” better.

#### 2.2.6 Internal diversity metrics and computer vision

Generative models in computer vision are also considering internal diversity metrics. For example, [17, section 4] introduced the *Inception score*, to assess both the quality and internal diversity of generated images. Other metrics are being considered and evaluated in the literature [18]. For future work, it will be interesting to build analogous metrics for molecule generation.

#### 2.2.7 Beyond fingerprints

Our definition of internal diversity depends on Morgan fingerprints, which are hand-crafted features that do not always capture the notion of chemical distance [19]. It would be better to use automatically learned features, molecule vector representations, analogous to word embeddings used in Natural Language Processing [20]. There is some work in this direction [21].

## 3 Generative Models

### 3.1 Reinforcement Learning

As in the case of RL considered in [8], and as in [22, p. 4], the generator *G_θ_* is a LSTM Recurrent Neural Network [23] parameterized by *θ*.

*G_θ_* maps the input embedding representations into a sequence of hidden states. Moreover, a softmax output layer maps the hidden states into the output token distribution. *G_θ_* generates SMILES (Simplified Molecular-Input Line-Entry System) sequences of length *T* (eventually padded with ”_” characters), denoted by:

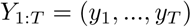

Let *R*(*Y*_1:*T*_) be the reward function.

- For the case of dopamine D2 activity, we take:

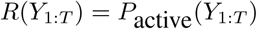

where *P*_active_(*Y*_1:*T*_)is the probability for *Y*_1:*T*_ to be D2-active. This probability is given by the predictive model made in [7]^[1]^, and available online at https://github.com/MarcusOlivecrona/REINVENT/releases
- For the case of druglikeness, we take:

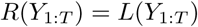

where *L*(*Y*_1:*T*_) is the druglikeness of *Y*_1:*T*_.

The generator *G_θ_* is viewed as a Reinforcement Learning agent: its state *s_t_* is the currently produced sequence of characters *Y*_1:*t*_, and its action *a* is the next character *y_t_*_+1_, which is selected in the alphabet *Y*. The agent policy is: *G_θ_*(*y_t_*_+1_*|Y*_1:*t*_). It corresponds to the probability to choose *y_t_*_+1_ given previous characters *Y*_1:*t*_.

Let *Q*(*s, a*) be the action-value function. It is the expected reward at state *s* for taking action *a* and for following the policy *G_θ_*, in order to complete the rest of the sequence. We maximize its expected long-term reward:

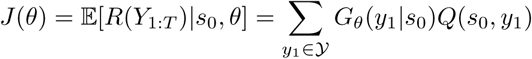

For any full sequence *Y*_1:*T*_, we have:

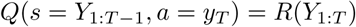

For *t < T*, in order to calculate the expected reward *Q* for *Y*_1:*t*_, we perform a *N*-time Monte Carlo search with the rollout policy *G_θ_*, represented as:

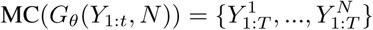

where 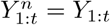 and 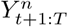 is randomly sampled via the policy *G_θ_*.

For *t < T*, *Q* is given by:

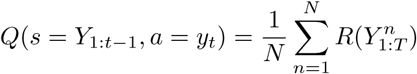

### 3.2 Objective-Reinforced Generative Adversarial Network (ORGAN)

To obtain an ORGAN, [8] brings a Character-Aware Neural Language Model [24] *D_ϕ_* parameterized by *ϕ*. Basically, *D_ϕ_* is a Convolutional Neural Network (CNN) whose output is given to a LSTM. *D_ϕ_* is fed with both training data and data generated by *G_θ_*. It plays the role of a *discriminator*, to distinguish between the two: for a SMILES *Y*_1:*T*_, the output *D_ϕ_*(*Y*_1:*T*_) is the probability that *Y*_1:*T*_belongs to the training data.

For the case of dopamine D2-activity, the reward function becomes:

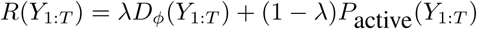

and for the case of druglikeness:

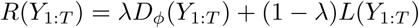

where *λ ∈* [0, 1] is a hyper-parameter. For *λ* = 0, we get back the RL case, and for *λ* = 1, we obtain a Sequential Generative Adversarial Network (SeqGAN) [22].

The networks *G_θ_* and *D_ϕ_* are trained adversarially [25, 26], such that the loss function for *D_ϕ_* to minimize is given by:

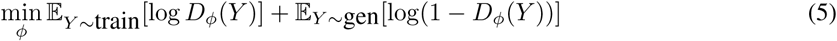

## 4 Experiments

### 4.1 Dopamine D2 activity

As in [8], we pre-train the models 240 steps with Maximum Likelihood Estimation (MLE), on a random subset of **15k molecules from the ZINC database** of 35 million commercially-available compounds for virtual screening, used in drug discovery [27]. Then we further train the models with RL and ORGAN respectively, for 30 and 60 steps more.

60 steps are enough to see how internal diversity drops. Training for longer increases the proportion of Valid SMILES, but it does not increase internal diversity of the desired molecules. See the second experiment. 30 steps are halfway between 0 and 60 steps, it allows to show the evolution.

Here are 5 samples from ZINC database (structures in figure 3):

**Figure 3:**
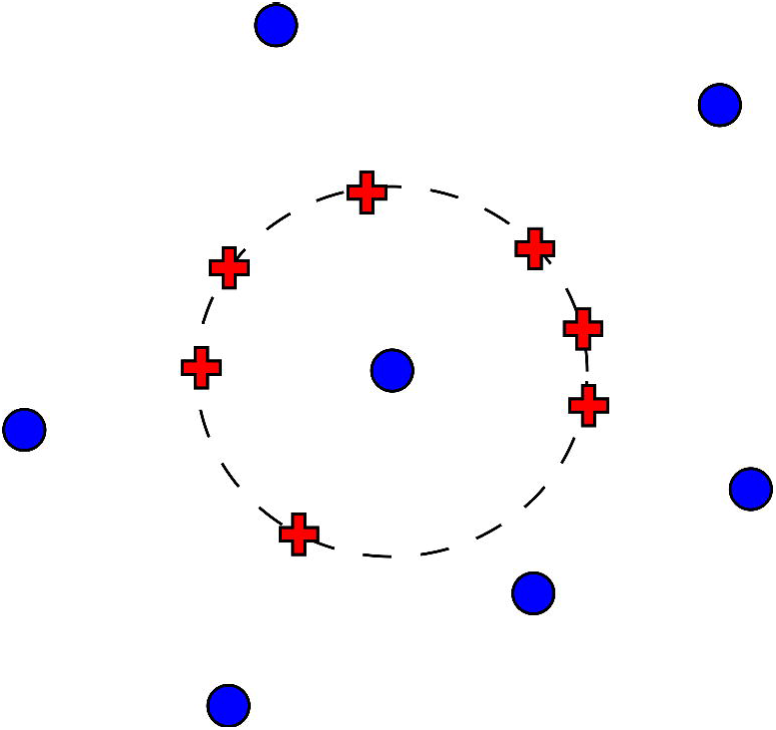
ZINC 15 structures.

COc1ccc(OC)c([C@@H]2CCCN2CN2C(=O)c3cccc(N)c3C2=O)c1

COc1cccc([C@@H](CNC(=O)Nc2ccc([N+](=O)[O-])cc2)[NH+](C)C)c1

COc1cccc(OC(=O)c2cc3ccccc3oc2=O)c1

CCCCC(=O)Nc1ccccc1NS(=O)(=O)c1ccc(Cl)cc1

COc1cc(CC[NH2+][C@H](C)c2cccnc2)ccc1F

In table 1, we show the proportion of valid SMILES output (Prop. Valid SMILES), the average probability of activity on dopamine D2 (Avg. *P_a_*), the average internal diversity (Avg. int. div.), the proportion of molecules with probability of activity greater than 0.8 (Prop. *P_a_ >* 0.8), and most importantly, the average internal diversity among samples with probability of activity greater than 0.8. That’s the most important column, because it is related with our open problem.

The averages are computed over the set of valid SMILES, whereas the proportions are computed over all the generated SMILES (both valid and invalid).

We compute those quantities for the model after the first 240 training epochs with Maximum Likelihood Estimation (MLE), a D2-active set of 8324 molecules from ExCAPE-DB [28] (which is essentially the training set of the SVM classifier in [7]) (DRD2), for the output of the Reinforcement Learning model after 30 steps (RL 30) and 60 steps (RL 60), and for the output of ORGAN with *λ* = 0.04 after 30 steps and 60 steps (ORGAN-0.04 30, ORGAN-0.04 60) and for *λ* = 0.5 after 60 steps (ORGAN-0.5 60). All those outputs have 32k samples.

The most interesting case is RL after 30 steps. In this case, increasing the probability of D2 activity is contradictory with keeping diversity. After 30 steps, internal diversity is even higher than the DRD2 diversity baseline.

However, when we only keep the molecules of interest, with *P_a_ >* 0.8, internal diversity dramatically drops to vanishingly small levels. Note that even in these cases, generated SMILES remain distinct from each other.

For ORGAN-0.04, results are mostly analogous to RL. Note that at 30 steps, diversity for *P_a_ >* 0.8 is 2 orders of magnitude better than RL 30. However, it still remains one order of magnitude lower than the DRD2 baseline, and at 60 steps, diversity has dropped to levels similar with RL.

For ORGAN-0.5, learning the D2 property still did not start after 60 steps. The situation is analogous to the SeqGAN case (*λ* = 1) described in [8]: high diversity, but no learning of the objective. In particular, that’s why the internal diversity for *P_a_ >* 0.8 is indetectable: there are only 6 samples satisfying the desired property, among 32k.

The intermediate cases between *λ* = 0.04 and *λ* = 0.5 are analogous to either of them. It is hard to situate the tipping point, between the cases where training is just slow, and where training will never take off.

Note that external diversity can be high, despite that internal diversity can be vanishingly small.

Here are the SMILES (structures in figure 4) of 10 samples for ORGAN with *λ* = 0.04 after 30 steps, selected such that *P_a_ >* 0.8 (most diverse case):

**Figure 4:**
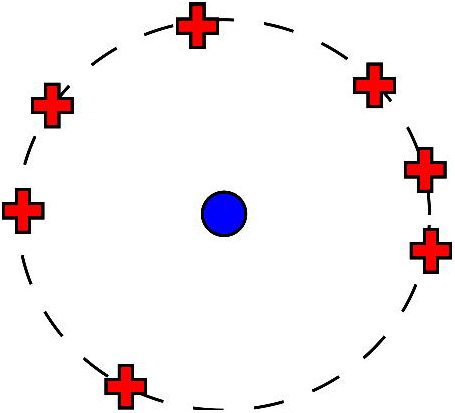
Generated structures for DRD2.

CCOCCNC[C@H]1CCCN1CCc1ccsc1

CCCOC[C@H]1Cc2ccccc21

CC[C@H]1CCNCOc2ccccc21

CC[C@H]1CCN(CCc2ccccc2)c1

CCCO[C@@H]1CCN(C)Cc2ccccc21

CCC[C@@H]1CCC[NH+]1CC[C@H]1CCCn1

CC[C@@H]1CCN(CCc2ccccc2)c1

CC[C@H]1CCN(Cc2ccccc2)c1

CCOC1CCN(CCCNCCCc2ccccc2)c1

CCCN1CCO[C@H]1C[C@@H]1CCOc2ccccc21

### 4.2 Druglikeness

Here, we use the experimental data from [8], made available on their Github: https://github.com/gablg1/ORGAN [8] pre-trains the models 240 epochs with Maximum Likelihood Estimation (MLE), on a random subset of 15k molecules from the ZINC database of 35 million commercially-available compounds for virtual screening, used in drug discovery [27]. Then [8] further trains the models with RL and ORGAN respectively, for 200 steps.

Table 2 shows the proportion of valid SMILES output (prop. Valid SMILES), average druglikeness (Avg. *L*), the average internal diversity (Avg. int. div.), the proportion of molecules with druglikeness greater than 0.8 (Prop. *L >* 0.8), and most importantly, the average internal diversity among samples with druglikeness greater than 0.8. Again, that’s the most important column, because it is related with our challenge.

**Table 2.** Experimental results for Druglikeness *L*

| | Prop. Valid SMILES | Avg. $L$ | Avg. int. div. | Prop. $L > 0.8$ | Avg. ext. div. for $L > 0.8$ | Avg. int. div. for $L > 0.8$ |
| --- | --- | --- | --- | --- | --- | --- |
| MLE | 0.290406 | 0.835164 | 0.393832 | 0.668891 | 0.347230 | 0.276438 |
| ZINC | 1 | 0.661094 | 0.331222 | 0.020133 | 0.235352 | 0.025986 |
| RL 200 | 0.975625 | 0.917358 | 0 | 0.974844 | 0.262789 | 0 |
| ORGAN-0.8 200 | 0.943906 | 0.906885 | 0.000151 | 0.940625 | 0.602854 | 0.000150 |

Again, the total averages are computed over the set of valid SMILES, whereas the proportions are computed over all the generated SMILES (both valid and invalid).

Those quantities are computed for the model after the first 240 training epochs with Maximum Likelihood Estimation (MLE), for the training set ZINC of 15k molecules (ZINC), which serves as a baseline, for the output of the Reinforcement Learning model after 200 steps (RL 200) and for the output of ORGAN with *λ* = 0.8 after 200 steps (ORGAN 200). *λ* = 0.8 was the highest parameter in experiments in [8]. Those outputs have 6400 samples.

Results show that ORGAN indeed improves over RL, since it is able to raise internal diversity to detectable levels. However, ORGAN diversity still remains 2 orders of magnitudes lower than ZINC diversity when *L >* 0.8. ORGAN diversity also remains 3 orders of magnitude lower than the total diversity of ZINC, which corresponds to the level of internal diversity to which most eyes are used to. We conclude that for our limited setting (small datasets...), both RL and ORGAN for *λ* = 0.8 fail to generate internally diverse molecules for this property.

Note again that external diversity can be very high, despite that internal diversity can be vanishingly small.

Note also that for ZINC, which is the training set, external and internal diversities for *L >* 0.8 are still different, because external diversity is taken over all training molecules, whereas internal diversity is taken over the small fraction (2 %) of them that have *L >* 0.8.

Here are 10 SMILES samples (structures in figure 5) from ORGAN for *λ* = 0.8 and 200 steps:

Cc1ccccc1CCSc1ccccc1C

COCCc1ccccc1CCCCCCSC

CCCCCCn1cccc1CCCSC

COCc1ccccc1CCCc1ccccc1CC

CCCC(=O)CCCc1ccccc1CCCc1ccccc1C

COCCNC(=O)CCc1ccccc1

CCOC(=O)CCCCc1ccccc1CCN(C)CCCCc1ccccc1C

COCCCC(=O)CC(C)CCCCc1ccccc1C

CCCC(=O)CSCCC(=O)CCCCC(=O)OC

CCC(=O)COCCCCCCC(=O)CCCSC

## 5 Conclusion and additional future work

The conclusion of this mini-benchmark is that for small training datasets, small architectures, and the specific hyperparameters tested, both RL and ORGAN fail to match observed internal diversity for desired molecules, although ORGAN is slightly better than RL. Future work about the diversity metrics was already discussed in subsection 2.2. There is also future work for a more comprehensive benchmark, with larger datasets, with larger and more models (like the recent [29]), and with testing various hyperparameters. Finally, there is also work for better ORGAN training. For this point, two distinct problems can be considered:

- The perfect discriminator problem in adversarial training
- The imbalance between different objectives in Reinforcement Learning

### 5.1 The perfect discriminator problem

In ORGAN training, the discriminator *D_ϕ_* quickly becomes perfect: it perfectly distinguishes between training data and generated data. In general, this situation is not very good for adversarial learning [30]. Here, the discriminator still teaches something to the generator. On average, according to the discriminator, the probability for a generated sample to belong to the training set still remains far from 0, although always smaller than 0.5. This probability is transmitted to the generator through the reward function.

However, not being able to ‘fool’ the discriminator, even in the SeqGAN case of *λ* = 1 (without any other objective), shows generator weakness: it shows inability to reproduce a plain druglike dataset like ZINC. Training a SeqGAN properly should be a first step towards improving ORGAN.

To achieve this, it might be possible to take a larger generator, to replace the discriminator loss in equation (5) with another function (like CramerGAN [31]), and to use one-sided label smoothing [17, p.4].

The discriminator might also overfit training data. Taking a larger training set could help, we took 15k samples here (less than 1MB), and this is small compared with training sets in Natural Language Processing. On the other hand, datasets in drug discovery rarely exceed 10k molecules, and therefore, it could also be interesting to look in the direction of low-data predictive neural networks [32].

Once adversarial training is stabilized, it might be interesting to replace all classifiers in the reward function with discriminators adversarially trained on different datasets. Various desired properties might be instilled into generated molecules with multiple discriminators. This might better transmit the chemical diversity present in the various training sets.

### 5.2 Imbalance in multi-objective RL

The main issue is the imbalance between the various objectives in the reward function, a problem occurring also in RL. Multi-objective reinforcement learning is a broad topic (for a survey, see [33]).

A problem here is that with a weighted sum, the agent always focuses on the easiest objective, and ignores harder ones. Moreover, the relative difficulty between objectives evolves over time. For example, the average probability of D2 activity initially grows exponentially, and so this growth is small when this probability is near 0.

Using time-varying adaptive weights might help. Moreover, those weights might not necessarily be linear: For example, the reward function can be of the form (*x^λ^* + *y^λ^*)^1*/λ*^, which converges towards min(*x, y*) as *λ → −∞*. Using an objective function of the form min(*x, y*) focuses the generator on the hard objective (but in our experiments, due to the perfect discriminator problem, it did not work).

Morever, in the reward function, a penalty can be introduced for newly generated molecules that are too similar with the generated molecules already having the desired properties.

In any case, the (varying) relative weights between different objectives must be determined automatically, and not through guesswork. In a drug discovery setting, a molecule must simultaneously satisfy a large number of objectives. For example, for an antipsychotic drug, it is not enough to be active against D2. The molecule must also pass toxicity and druglikeness tests. Moreover, to avoid side-effects, the molecule must not be active with D3, D4, serotonin, or histamine. That’s a lot of objectives to include in the reward function.

## Availability of data and material

All code and data are available at: https://github.com/mostafachatillon/ChemGAN-challenge

## Acknowledgements

Computations were performed with 2 GPUs Nvidia Tesla M60, available at Microsoft Azure Free Trial.

---

Figure 5: **Generated structures for druglikeness**.

## Additional Material

Additional file 1 — trivialmodel.jpeg Figure 1

Additional file 2 — nontrivialmodel.jpeg Figure 2

Additional file 3 — zinc15.jpg Figure 3

Additional file 4 — drd2 generated.jpg Figure 4

Additional file 5 — druglikeness.jpg Figure 5

Code and data — https://github.com/mostafachatillon/ChemGAN-challenge

## Footnotes

[1] This reward function is slightly different than the function in [7], which is: *−*1 + 2 *× P*active.

## References

1. DiMasi, J.A., Grabowski, H.G., Hansen, R.W.: Innovation in the pharmaceutical industry: new estimates of r&d costs. Journal of health economics 47, 20–33 (2016)

2. Gómez-Bombarelli, R., Duvenaud, D., Hernández-Lobato, J.M., Aguilera-Iparraguirre, J., Hirzel, T.D., Adams, R.P., Aspuru-Guzik, A.: Automatic chemical design using a data-driven continuous representation of molecules. arXiv preprint arXiv:1610.02415 (2016)

3. Kadurin, A., Aliper, A., Kazennov, A., Mamoshina, P., Vanhaelen, Q., Khrabrov, K., Zhavoronkov, A.: The cornucopia of meaningful leads: Applying deep adversarial autoencoders for new molecule development in oncology. Oncotarget 8(7), 10883 (2017)

4. Kadurin, A., Nikolenko, S., Khrabrov, K., Aliper, A., Zhavoronkov, A.: drugan: an advanced generative adversarial autoencoder model for de-novo generation of new molecules with desired molecular properties in silico. Molecular Pharmaceutics (2017)

5. Jaques, N., Gu, S., Bahdanau, D., Hernández-Lobato, J.M., Turner, R.E., Eck, D.: Sequence tutor: Conservative fine-tuning of sequence generation models with kl-control. In: International Conference on Machine Learning, pp. 1645–1654 (2017)

6. Segler, M.H., Kogej, T., Tyrchan, C., Waller, M.P.: Generating focussed molecule libraries for drug discovery with recurrent neural networks. arXiv preprint arXiv:1701.01329 (2017)

7. Olivecrona, M., Blaschke, T., Engkvist, O., Chen, H.: Molecular de novo design through deep reinforcement learning. Journal of Cheminformatics 9(1), 48 (2017)

8. Guimaraes, G.L., Sanchez-Lengeling, B., Farias, P.L.C., Aspuru-Guzik, A.: Objective-reinforced generative adversarial networks (organ) for sequence generation models. arXiv preprint arXiv:1705.10843 (2017)

9. Benjamin, S.-L., Carlos, O., L., G.G., Alan, A.-G.: Optimizing distributions over molecular space. an objective-reinforced generative adversarial network for inverse-design chemistry (organic). ChemRxiv Preprint https://doi.org/10.26434/chemrxiv.5309668.v3 (2017)

10. Bickerton, G.R., Paolini, G.V., Besnard, J., Muresan, S., Hopkins, A.L.: Quantifying the chemical beauty of drugs. Nature chemistry 4(2), 90–98 (2012)

11. Ertl, P., Schuffenhauer, A.: Estimation of synthetic accessibility score of drug-like molecules based on molecular complexity and fragment contributions. Journal of cheminformatics 1(1), 8 (2009)

12. Rogers, D., Hahn, M.: Extended-connectivity fingerprints. Journal of chemical information and modeling 50(5), 742–754 (2010)

13. Landrum, G.: Rdkit: Open-source cheminformatics. http://www.rdkit.org (2017)

14. Gupta, A., Mü ller, A.T., Huisman, B.J., Fuchs, J.A., Schneider, P., Schneider, G.: Generative recurrent networks for de novo drug design. Molecular informatics (2017)

15. Merk, D., Friedrich, L., Grisoni, F., Schneider, G.: De novo design of bioactive small molecules by artificial intelligence. Molecular informatics (2018)

16. Wattenberg, M., Viégas, F., Johnson, I.: How to use t-sne effectively. Distill (2016). doi:10.23915/distill.00002

17. Salimans, T., Goodfellow, I., Zaremba, W., Cheung, V., Radford, A., Chen, X.: Improved techniques for training gans. In: Advances in Neural Information Processing Systems, pp. 2234–2242 (2016)

18. Anonymous: An empirical study on evaluation metrics of generative adversarial networks. International Conference on Learning Representations (2018)

19. Giordanetto, F., Boström, J., Tyrchan, C.: Follow-on drugs: How far should chemists look? Drug discovery today 16(15), 722–732 (2011)

20. Mikolov, T., Sutskever, I., Chen, K., Corrado, G.S., Dean, J.: Distributed representations of words and phrases and their compositionality. In: Advances in Neural Information Processing Systems, pp. 3111–3119 (2013)

21. Wu, Z., Ramsundar, B., Feinberg, E.N., Gomes, J., Geniesse, C., Pappu, A.S., Leswing, K., Pande, V.: Moleculenet: A benchmark for molecular machine learning. arXiv preprint arXiv:1703.00564 (2017)

22. Yu, L., Zhang, W., Wang, J., Yu, Y.: Seqgan: sequence generative adversarial nets with policy gradient. In: AAAI-17: Thirty-First AAAI Conference on Artificial Intelligence, vol. 31 (2017). Association for the Advancement of Artificial Intelligence

23. Hochreiter, S., Schmidhuber, J.: Long short-term memory. Neural computation 9(8), 1735–1780 (1997)

24. Kim, Y., Jernite, Y., Sontag, D., Rush, A.M.: Character-aware neural language models. In: AAAI, pp. 2741–2749 (2016)

25. Schmidhuber, J.: Learning factorial codes by predictability minimization. Neural computation 4(6), 863–879 (1992)

26. Goodfellow, I., Pouget-Abadie, J., Mirza, M., Xu, B., Warde-Farley, D., Ozair, S., Courville, A., Bengio, Y.: Generative adversarial nets. In: Advances in Neural Information Processing Systems, pp. 2672–2680 (2014)

27. Sterling, T., Irwin, J.J.: Zinc 15—ligand discovery for everyone. J. Chem. Inf. Model 55(11), 2324–2337 (2015)

28. Sun, J., Jeliazkova, N., Chupakin, V., Golib-Dzib, J.-F., Engkvist, O., Carlsson, L., Wegner, J., Ceulemans, H., Georgiev, I., Jeliazkov, V., et al.: Excape-db: an integrated large scale dataset facilitating big data analysis in chemogenomics. Journal of Cheminformatics 9(1), 17 (2017)

29. Dai, H., Tian, Y., Dai, B., Skiena, S., Song, L.: Syntax-directed variational autoencoder for molecule generation. NIPS Workshop (2017)

30. Arjovsky, M., Bottou, L.: Towards principled methods for training generative adversarial networks. arXiv preprint arXiv:1701.04862 (2017)

31. Bellemare, M.G., Danihelka, I., Dabney, W., Mohamed, S., Lakshminarayanan, B., Hoyer, S., Munos, R.: The cramer distance as a solution to biased wasserstein gradients. arXiv preprint arXiv:1705.10743 (2017)

32. Altae-Tran, H., Ramsundar, B., Pappu, A.S., Pande, V.: Low data drug discovery with one-shot learning. ACS central science 3(4), 283–293 (2017)

33. Roijers, D.M., Vamplew, P., Whiteson, S., Dazeley, R.: A survey of multi-objective sequential decision-making. Journal of Artificial Intelligence Research 48, 67–113 (2013)

